# Induction of Chk2 signaling in the germarium is sufficient to cause oogenesis arrest in *Drosophila*

**DOI:** 10.1101/611798

**Authors:** Zeljko Durdevic, Anne Ephrussi

## Abstract

The conserved RNA helicase Vasa is required for germ cell development in many organisms. It is established that in *Drosophila* loss of piRNA pathway components, including Vasa, causes Chk2-dependent oogenesis arrest, however the stage at which Chk2-signaling is triggered was unknown. We found that absence of Vasa during the germarial stages arrests oogenesis due to Chk2 activation. Importantly, once induced in the germarium, Chk2-mediated arrest of oogenesis cannot be overcome by restoration of Vasa to the arrested egg-chambers. We conclude that Vasa activity specifically in the germarium is essential for germ cell lineage development.

---

Development of the *Drosophila* female gonad begins during the third larval instar with the formation of 16-25 somatic niches that will give rise to the future germaria (Panchal et al., 2017). Each germarium hosts germ line stem cells (GSCs) that produce the germ cell lineage (Wieschaus and Szabad, 1979). In adult females, germ cell development begins with the division of a GSC into a self-renewing stem cell and a differentiating daughter cell, the cystoblast (CB). The CB undergoes four rounds of mitosis with incomplete cytokinsesis, such that a stage 1 egg-chamber is ultimately composed of an oocyte and 15 nurse cells, surrounded by a layer of follicular epithelial cells (reviewed in Gilboa and Lehmann, 2004). A newly formed egg-chamber buds off from the germarium and joins a linear array of developing egg-chambers to form an ovariole. Each *Drosophila* ovary consists of 16-25 ovarioles, corresponding to the number of germaria formed in the third instar larva.

*Drosophila* oogenesis has been intensively studied and many genes found to regulate development of the germ cell lineage. Among the germline proteins essential for oogenesis is the conserved RNA helicase Vasa (Vas). Vas is expressed throughout oogenesis and localizes to the posterior pole of the oocyte and early embryo. In situations leading to absence of Vas from the oocyte posterior pole, germline and posterior patterning determinants fail to localize, germ (or pole) plasm does not form, and the resulting embryos lack posterior structures and primordial germ cells (Hay et al., 1990; Lasko and Ashburner, 1988, 1990). In contrast to late oogenesis and embryos, little is known about the role of Vas during early oogenic stages. In early oogenesis *vas* has been implicated in the translational control of *mei-p26* and in regulation of GSC mitotic chromosome condensation (Liu et al., 2009; Pek and Kai, 2011). As complete absence of *vas* triggers oogenesis arrest induced by *checkpoint kinase 2* (*Chk2*) (Durdevic et al., 2018; Lasko and Ashburner, 1990), the question remains at which stage Vas activity is critically required for oogenesis to complete.

To address the importance of Vas activity in early germ cell lineage development we took a genetic approach. We found that targeted exclusion of Vas from the germarium arrests oogenesis and reduces the number of egg-chamber-producing ovarioles. Furthermore, in the absence of Vas in the germarium, Chk2-signaling induces germ cell developmental arrest. Importantly, once induced in the germarium, Chk2-mediated arrest of oogenesis cannot be overcome by restoration of Vas to the arrested egg-chambers. Our data show that the activity of Vas RNA helicase early in oogenesis prevents activation of Chk2-signaling, ensuring sustained development of the germline component of the ovary.

## Materials and Methods

### Fly stocks and husbandry

The following *Drosophila* stocks were used: *w*^*1118*^; *vas*^*PD*^*/CyO* (*vas*^*1*^, Schüpbach and Wieschaus, 1986); *vas*^*D1*^*/CyO* (*vas*^*3*^, Tearle and Nusslein-Volhard, 1987, Lasko and Ashburner, 1990); *vas-Gal4* (gift of Jean-René Huynh); *matTub-Gal4-VP16* (gift of Stefano De Renzis); *GFP-vas*^*WT*^*/TM2, and GFP-vas*^*DQAD*^*/TM2* and *nos-Gal4-VP16/TM2* (Xiol et al., 2014); (Xiol et al., 2014); *P{TRiP.GL00020}attP2/TM3* (TRiPmnk, FBst0035152); *P{TRiP.GL00094}attP2* (TRiPw, FBst0035573); *P{Ubi-GFP.S65T}PAD1* (GFP, FBst0004888). All flies were kept at 25°C on standard *Drosophila* medium.

### Generation of transgenic flies and expression of the transgenes

The *vasa* transgene carrying the K295N substitution *(vas*^*GNT*^) was created by introducing the point mutation in wild-type *vasa* cDNA sequence by site directed mutagenesis using QuikChange II XL Site-Directed Mutagenesis Kit (Agilent). The transgene is subsequently cloned downstream of GFP into the pUASK10attB vector (gift of Beat Suter), from which the K10 sequence was removed. Transgenes were integrated into the attP transposable element insertion site of the landing site line VK33 (P (nos-phiC31\int.NLS)) X, PBac(y[+]-attP-3B)VK00033, gift of Hugo Bellen).

All transgenes are expressed using the GAL4/UAS-system. Gal4-drivers were under control of two promoters with distinct expression patterns: vas-Gal4 is expressed throughout oogenesis and matTub-Gal4 is excluded from the germarium (Supplementary Figure S1B).

### Fecundity and hatching assays

Virgin females of *w*^*1118*^ control and *vas*^*PD/D1*^ and *vas*^*D1/D1*^ genetic backgrounds with or without expressed transgenes were mated with *w*^*1118*^ males for 24h at 25°C. The crosses were then transferred to apple-juice agar plates, which were used to collect eggs at 24h intervals over 3 or 3-20 days. The number of laid eggs on each plate was counted and the plates were kept at 25°C for 24h and the number of hatched larvae was also counted (Supplementary Table S1 and S2). Experiments were performed in five independent replicates.

### Ovarian morphology and quantification of ovariole number

Ovaries were dissected from 3-day to 20-day old flies in PBS. To assess ovarian morphology, ovaries were directly imaged on Olympus SZX16 stereomicroscope. The length of the ovaries was measured using *Fiji* (Supplementary Table S3, S4 and S5). For determination of egg-chamber-producing ovariole number females were frozen and held at −20°C prior dissection. Ovaries were manually dissected under magnification in a drop of PBS. The ovarioles were gently separated from each other using wolfram needles. The ovariole number of each female was defined as a summary of the number of egg-chamber-containing ovarioles in the right and left ovary (Supplementary Table S6 and S7).

### Vasa localization analysis

For Vas localization, ovaries from wild-type flies (w^1118^) and *vas* mutants (*vas*^*PD/D1*^ and *vas*^*D1/D1*^) were fixed by incubation at 92°C for 5 min in preheated fixation buffer (0.4% NaCl, 0.3% Triton X-100 in PBS), followed by extraction in 1% Triton X-100 for 1h at room temperature (RT). Fixed ovaries were incubated with primary antibodies Vasa (rat; 1:500; (Tomancak et al., 1998)) and subsequently with secondary antibodies Alexa 647 conjugated donkey anti-rat IgG (1:1000; Jackson ImmunoResearch). Ovaries from flies expressing fusion proteins were fixed in 2% PFA and 0.01% Triton X-100 for 15 min at RT. Fixed ovaries were mounted on glass slides for examination of GFP fluorescence for the fusion proteins and Alexa 647 fluorescence for wild-type Vas using a Zeiss LSM 780 confocal microscope. Nuclei were visualised with DAPI.

### Cuticle preparation

To examine larval cuticles, eggs were allowed to develop fully for 24h at 25°C, dechorionated in bleach, and then transferred to a microscope slide bearing a drop of Hoyer’s medium mixed 1:1 with lactic acid. Cuticle preparations were heated at 65° overnight before examination using Zeiss Axiophot microscope. Number of counted larvae with or without abdomen is represented in Supplementary Table S8.

### Immunohistochemical staining of ovaries and embryos

Freshly hatched females were mated with wild-type males and kept for 2-3 days on yeast at 25°C prior to dissection. Ovaries were dissected in PBS and immediately fixed by incubation at 92°C for 5 min in preheated fixation buffer (0.4% NaCl, 0.3% Triton X-100 in PBS), followed by extraction in 1% Triton X-100 for 1h at room temperature (RT). Fixed ovaries were incubated with primary antibodies Aub (rabbit; 1:500; (Homolka et al., 2015)), Ago3 (mouse; 1:250; gift of Mikiko Siomi), Vasa (rat; 1:500; (Tomancak et al., 1998)). The following secondary antibodies were used: Alexa 488 conjugated goat anti-rabbit (1:1000; Invitrogen) and anti-mouse IgG (1:500; Invitrogen), Alexa 647 conjugated donkey anti-rat IgG (1:1000; Jackson ImmunoResearch). Nuclei were stained with DAPI.

For embryo staining, freshly hatched females were mated with wild-type males and fed with yeast for 2-3 days at 25°C prior to egg collection. Embryos (0-1h or 1-3h) were collected and dechorionated in 50% bleach, then fixed by incubation at 92°C for 30 sec in preheated fixation buffer (0.4% NaCl, 0.3% Triton X-100 in PBS), followed by devitellinization by rigorous shaking in a 1:1 mix of heptane and methanol. After washing in 0.1% Tween-20, embryos were either immediately incubated with primary antibodies against Aub (rabbit; 1:500; (Homolka et al., 2015)), Ago3 (mouse; 1:250; gift of Mikiko Siomi), Vasa (rat; 1:500; (Tomancak et al., 1998)), or stored in methanol at −20°C for staining later on. The following secondary antibodies were used: Alexa 488 conjugated goat anti-rabbit (1:1000; Invitrogen) and anti-mouse IgG (1:500; Invitrogen), Alexa 647 conjugated donkey anti-rat IgG (1:1000; Jackson ImmunoResearch). Nuclei were stained with DAPI.

The samples were observed using a Zeiss LSM 780 or Leica SP8 confocal microscope. Oocytes and embryos with Aub and Ago3 positive pole plasm were counted in three independent replicates (Supplementary Table S9 and S10).

### Protein extraction and western blotting

For whole protein lysates of ovaries, around 20 pairs of ovaries from 3-5-day old flies were homogenized in protein extraction buffer (25 mM Tris pH 8.0, 27.5 mM NaCl, 20 mM KCl, 25 mM sucrose, 10 mM EDTA, 10 mM EGTA, 1 mM DTT, 10% (v/v) glycerol, 0.5% NP40, 1% Triton X-100, 1x Protease inhibitor cocktail (Roche)). For whole protein embryo lysates, 0-2h or 1-3h old embryos were collected from apple-juice agar plates and homogenized in protein extraction buffer. Samples were incubated on ice for 10 min, followed by two centrifugations, each 15 min at 16.000 g. 50-100 μg of total protein extracts were solubilized in SDS-sample buffer by boiling at 95°C for 5 minutes and analysed by SDS polyacrylamide gel electrophoresis (4-12% NuPAGE gel; Invitrogen). Western blotting was performed using antibodies against Vasa (rat; 1:3000; (Tomancak et al., 1998)), Aub (rabbit; 1:1000; (Homolka et al., 2015), Ago3 (mouse; 1:500; gift of Mikiko Siomi), GFP (rabbit; 1:5000; Chemokine TP401), and Tub (mouse; 1:10000; Sigma T5168).

Quantification of relative protein expression levels was performed using ImageJ. A frame was placed around the most prominent band on the image and used as a reference to measure the mean gray value of all other protein bands, as well as the background. Next, the inverted value of the pixel density was calculated for all measurements by deducting the measured value from the maximal pixel value. The net value of target proteins and the loading control was calculated by deducting the inverted background from the inverted protein value. The ratio of the net value of the target protein and the corresponding loading control represents the relative expression level of the target protein. Fold-change was calculated as the ratio of the relative expression level of the target protein in the wild-type control over that of a specific sample.

### Protein immunoprecipitation and proteomic analysis

For protein immunoprecipitation, ovaries of 3-5 day-old *vas*^*PD/D1*^*; vas-Gal4>GFP-vas*^*WT*^*, vas*^*PD/D1*^*; vas-Gal4>GFP-vas*^*DQAD*^ and control *Act5C-Gal4/CyO; Ubq-GFP* flies were dissected in PBS and homogenized in protein extraction buffer (25 mM Tris pH 8.0, 27.5 mM NaCl, 20 mM KCl, 25 mM sucrose, 10 mM EDTA, 10 mM EGTA, 1 mM DTT, 10% (v/v) glycerol, 0.5% NP40, 1% Triton X-100, 1x Protease inhibitor cocktail (Roche)). After 10 min incubation on ice an equal volume of protein extraction buffer without detergents was added to the samples, and the samples were centrifuged twice for 15 min at 16,000 g. Immunoprecipitations were performed using GFP-trap magnetic agarose beads (ChromoTek) at 4°C for 1h on 20 mg of protein lysates. The beads were washed 5 times for 5 min at 4°C: 1x in protein extraction buffer, then 1x in high salt buffer (25 mM Tris pH 8.0, 1M NaCl, 0.5% NP40, 1% Triton X-100, 1x Protease inhibitor cocktail (Roche)), then in 1x medium salt buffer (25 mM Tris pH 8.0, 0.5M NaCl, 0.5% NP40, 1% Triton X-100, 1x Protease inhibitor cocktail (Roche)), then in 1x low salt buffer (25 mM Tris pH 8.0, 150mM NaCl, 0.5% NP40, 1% Triton X-100, 1x Protease inhibitor cocktail (Roche)) and finally in 1x low salt buffer without detergents (25 mM Tris pH 8.0, 150mM NaCl, 1x Protease inhibitor cocktail (Roche)). After washing, precipitated proteins were eluted from the magnetic beads in elution buffer (200mM glycine pH 2.5) and neutralized with 1/10 volume of 1M Tris base pH 10.4.

Proteomic experiments were performed as described in (Casabona et al., 2013). In brief, three samples were stacked in the SDS polyacrylamide gel and after Coomassie staining each lane was cut into 3 blocks, which were processed separately. After digestion with trypsin (Promega, sequencing grade), the resulting peptides were analyzed by LC-MS/MS (LTQ-Orbitrap Velos pro, Thermo Fisher Scientific). Peptides and proteins from each MS run were identified using Scaffold software and results for each lane were displayed. The selection criteria for the displayed proteins were: a minimum of 2 peptides per protein should be identified and the peptide Mascot score should be at least 20. The experiment was performed in two biological replicates and specific interaction partners were determined by statistical analysis of control and positive samples using extracted spectral counts. A protein was considered as a high confidence binding partner if its enrichment was equal to or above 2 and the *p*-value was ≤ 0.05 in positive IPs compared to controls. All proteins that showed enrichment equal to or above 2 but had a higher *p*-value were considered low confidence hits. *P*-values were computed using the web-based Quantitative Proteomics *p*-value Calculator (QPPC) (Chen et al., 2014) that applies a distribution-free permutation method based on simulation of the log(ratio). A pseudocount of 1 was used in all samples for proteins with no spectral counts. Un-weighted spectrum counts for both replicates and the results of the statistical analysis are provided in Supplementary Table S11.

### Fluorescent *in situ* RNA hybridization

FISH experiments were performed as described in (Gaspar et al., 2017). In brief, ovaries were dissected in PBS and immediately fixed in 2% PFA, 0.05 % Triton X-100 in PBS for 20 min at RT. After washing in PBT (PBS + 0.1% Triton X-100) samples were treated with 2 μg/mL proteinase K in PBT for 5 min and then were subjected to 95°C in PBS + 0.05% SDS for 5 min. Samples were pre-hybridized in 200 μL hybridization buffer (300 mM NaCl, 30 mM sodium citrate pH 7.0, 15 % ethylene carbonate, 1 mM EDTA, 50 μg/mL heparin, 100 μg/mL salmon sperm DNA, 1% Triton X-100) for 10 min at 42°C. Fluorescently labeled oligonucleotides (12.5–25 nM) were pre-warmed in hybridization buffer and added to the samples. Hybridization was allowed to proceed for 2 h at 42°C. Samples were washed 3 times for 10 min at 42°C in pre-warmed buffers (1x hybridization buffer, then 1x hybridization buffer:PBT 1:1 mixture, and then 1x PBT). The final washing step was performed in pre-warmed PBT at RT for 10 min. The samples were mounted in 80% 2,2-thiodiethanol in PBS and analyzed on a Leica SP8 confocal microscope.

### Labeling of DNA oligonucleotides for fluorescent *in situ* RNA hybridization

Labeling of the oligonucleotides was performed as described in (Gaspar et al., 2017). Briefly, non-overlapping arrays of 18-22 nt long DNA oligonucleotides complementary to *mnk* (Supplementary Table S12) were selected using the *smFISHprobe_finder.R* script (Gaspar et al., 2017). An equimolar mixture of oligos for a given RNA was fluorescently labelled with Alexa 565− or Alexa 633-labeled ddUTP using terminal deoxynucleotidyl transferase. After ethanol precipitation and washing with 80% ethanol, fluorescently labeled oligonucleotides were reconstituted with nuclease-free water.

### RNA extraction and quantitative PCR analysis

Total RNA was extracted from ovaries of 3-day old flies using Trizol reagent (Thermofisher). For first-strand cDNA synthesis, RNA was reverse transcribed using the QuantiTect Reverse Transcription Kit (QIAGEN). Quantitative PCR was performed on a StepOne Real-Time PCR System (Thermofisher) using SYBR Green PCR Master Mix (Thermofisher). Experiments were performed in biological triplicates with technical triplicates. Relative RNA levels were calculated by the 2^-ΔΔCT^ method (Livak and Schmittgen, 2001) and normalized to *rp49* mRNA levels and normalized to respective RNA levels from *w*^*1118*^ flies. Sequences of primers used for qPCR reaction are presented in Supplementary Table S12.

## Results

### Vasa helicase activity is required for *Drosophila* oogenesis

To investigate effects of Vas helicase activity on germ cell lineage development, we used the UAS/GAL4 system to manipulate the expression of wild-type Vas and of two Vas helicase mutants. The Vas mutant proteins contain amino acid substitutions that affect helicase activity at different points in the enzymatic process: the K295N (GKT→GNT) substitution hinders ATP and RNA binding and locks the helicase in an open conformation, whereas the E400Q (DEAD→DQAD) mutation prevents release of the ATP-hydrolysis products and locks the helicase in a closed conformation (Xiol et al., 2014) (Supplementary Figure S1A). To monitor expression and localization of the different Vas proteins, we fused them with GFP (GFP-Vas^DQAD^, GFP-Vas^GNT^, GFP-Vas^WT^).

We tested the ability of the proteins to provide Vas function and suppress the oogenesis arrest displayed by *vas*^*D1/D1*^ mutants and rescue the abdominal defects (posterior group phenotype) of embryos produced by the hypomorphic *vas*^*PD/D1*^ females. We assessed ovarian morphology and quantified ovary length as a measure of oogenesis rescue (Supplementary Figure S1C). The GFP-Vas^WT^ transgene fully restored oogenesis to the loss of function *vas*^*D1/D1*^ flies, whereas the helicase mutants GFP-Vas^DQAD^ and GFP-Vas^GNT^ did not (Figure 1A and Supplementary Figure S1D). Furthermore, Vas function provided by the GFP-Vas^WT^ transgene promoted abdomen formation in around 50% of embryos produced by both *vas*^*D1/D1*^ and *vas*^*PD/D1*^ females (Figure 1B-C). In contrast, the embryos produced by helicase inactive GFP-Vas^DQAD^ and GFP-Vas^GNT^ expressing *vas*^*PD/D1*^ females had a strong posterior group phenotype and did not hatch (Figure 1B-C). These results suggest that the helicase activity of Vas is required for oogenesis and embryogenesis.

**Figure 1.**
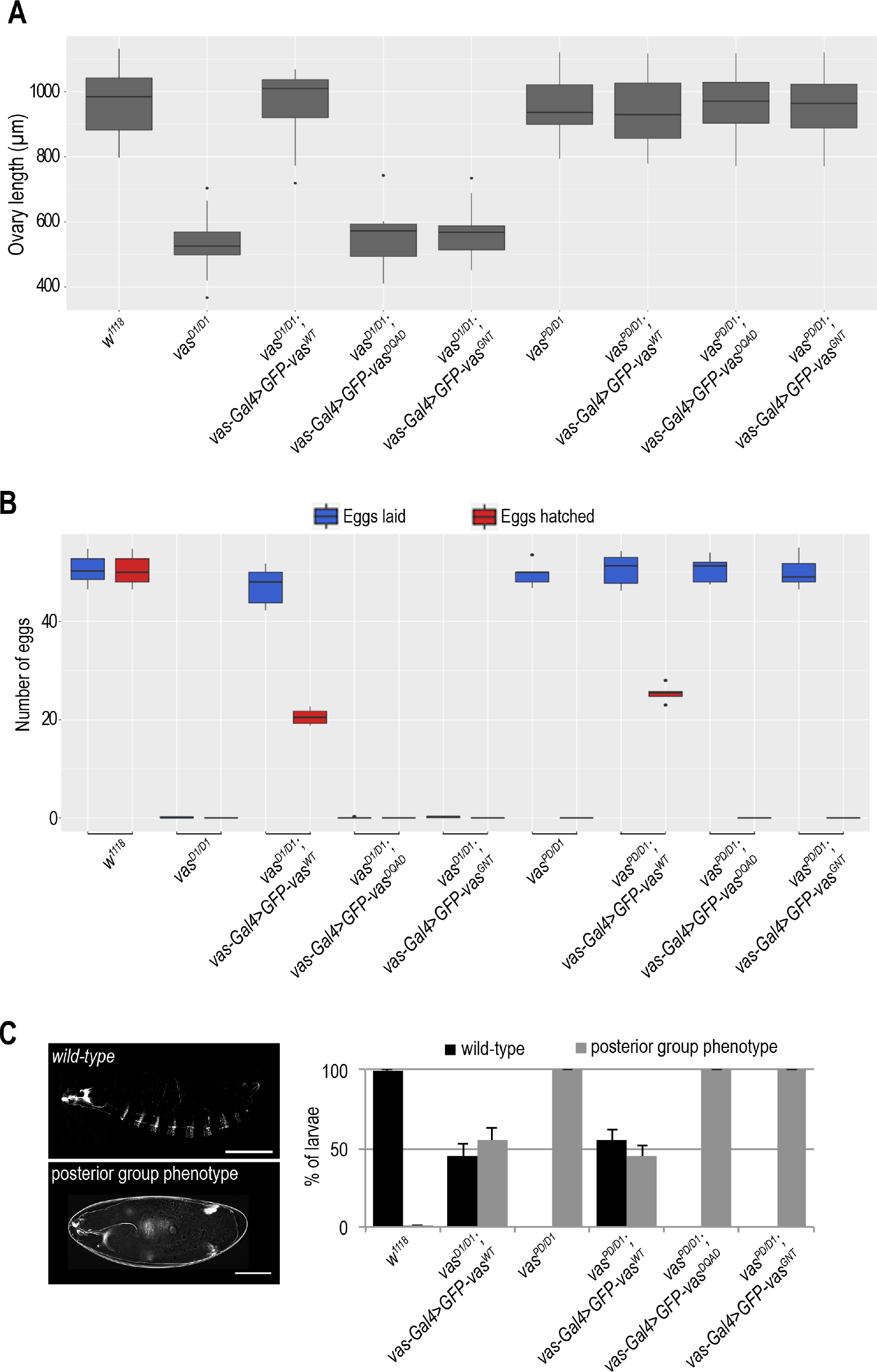
Vasa helicase activity is essential for germline and embryo development. A) Box plot representing length of ovaries of wild-type (*w*^*1118*^*), vas*^*D1/D1*^, *vas*^*D1/D1*^*; vas-Gal4>GFP-Vas*^*WT*^, *vas*^*D1/D1*^*; vas-Gal4>GFP-Vas*^*DQAD*^, *vas*^*D1/D1*^*; vas-Gal4>GFP-Vas*^*GNT*^, *vas*^*PD/D1*^, *vas*^*PD/D1*^*; vas-Gal4>GFP-Vas*^*WT*^, *vas*^*PD/D1*^*; vas-Gal4>GFP-Vas*^*DQAD*^ and *vas*^*PD/D1*^*; vas-Gal4>GFP-Vas*^*GNT*^ flies. The measurements were performed on 10 flies (n=10) of each genotype. B) Box plot representing the number of eggs laid and hatched from wild-type (*w*^*1118*^*), vas*^*D1/D1*^, *vas*^*D1/D1*^*; vas-Gal4>GFP-Vas*^*WT*^, *vas*^*D1/D1*^*; vas-Gal4>GFP-Vas*^*DQAD*^, *vas*^*D1/D1*^*; vas-Gal4>GFP-Vas*^*GNT*^, *vas*^*PD/D1*^, *vas*^*PD/D1*^*; vas-Gal4>GFP-Vas*^*WT*^, *vas*^*PD/D1*^*; vas-Gal4>GFP-Vas*^*DQAD*^ and *vas*^*PD/D1*^*; vas-Gal4>GFP-Vas*^*GNT*^ flies. Five independent replicates of the experiment were performed. C) Larval cuticle phenotypes observed in wild-type (*w*^*1118*^*), vas*^*D1/D1*^*; vas-Gal4>GFP-Vas*^*WT*^, in *vas*^*PD/D1*^*, vas*^*PD/D1*^*; vas-Gal4>GFP-Vas*^*WT*^, *vas*^*PD/D1*^*; vas-Gal4>GFP-Vas*^*DQAD*^ and *vas*^*PD/D1*^*; vas-Gal4>GFP-Vas*^*GNT*^ flies. Scale bars indicate 500 μm (larva) and 100 μm (un-hatched larva).

### Localization of PIWI proteins is affected by Vasa’s helicase activity

Localization of Vas in the egg-chamber is independent of RNA-binding and helicase activity (Dehghani and Lasko, 2016; Liang et al., 1994). We analyzed whether the E400Q and K295N mutations (Supplementary Figure S1A), which impair Vas helicase activity differently, locking the enzyme in a closed or in an open conformation, respectively, affect localization of the protein. Additionally, as Aub and Ago3 co-localize with Vas (Liang et al., 1994; Malone et al., 2009), we tested if the localization of these two PIWI proteins in the egg-chamber and embryo is affected by Vas helicase mutations.

Localization to nurse cell nuage was impaired in the case of GFP-Vas^DQAD^ (closed conformation) (Xiol et al., 2014), whereas GFP-Vas^WT^ and GFP-Vas^GNT^ (open conformation) showed wild-type localization (Figure 2A-B). This was true for the localization of Aub and Ago3 (Supplementary Figure S2A-B and S3A). These findings indicate that an open helicase conformation of the Vas is required for its correct localization, as well as for the localization of Aub and Ago3, whereas helicase activity per se is not.

**Figure 2:**
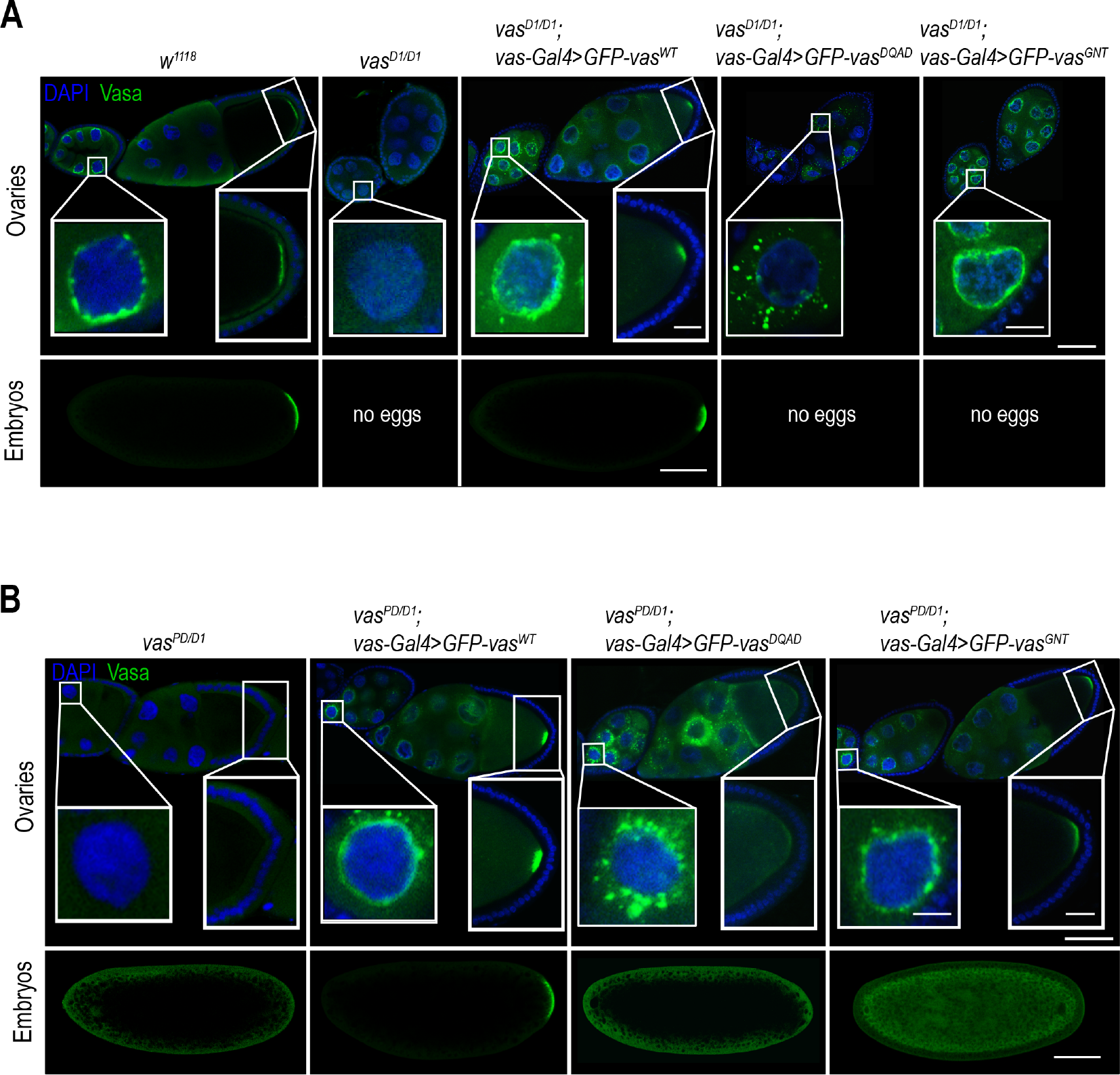
Helicase activity is required for Vasa localization at the posterior pole of the embryo. A) Localization of Vasa in egg-chambers (upper panels) and embryos (bottom panels) of wild-type (*w*^*1118*^*), vas*^*D1/D1*^, *vas*^*D1/D1*^*; vas-Gal4>GFP-Vas*^*WT*^, *vas*^*D1/D1*^*; vas-Gal4>GFP-Vas*^*DQAD*^ and *vas*^*D1/D1*^*; vas-Gal4>GFP-Vas*^*GNT*^ flies. Scale bars indicate 50 μm (egg-chamber), 10 μm (nuage and pole plasm) and 100 μm (embryo). B) Localization of Vasa in egg-chambers (upper panels) and embryos (bottom panels) of *vas*^*PD/D1*^, *vas*^*PD/D1*^*; vas-Gal4>GFP-Vas*^*WT*^, *vas*^*PD/D1*^*; vas-Gal4>GFP-Vas*^*DQAD*^ and *vas*^*PD/D1*^*; vas-Gal4>GFP-Vas*^*GNT*^ flies. Scale bars indicate 50 μm (egg-chamber), 10 μm (nuage and pole plasm) and 100 μm (embryo).

In oocytes and embryos, GFP-Vas^WT^ showed a wild-type localization at the posterior pole (Figure 2A-B), whereas GFP-Vas^DQAD^ was not detected. And, although we could detect GFP-Vas^GNT^ at the posterior pole of the oocyte and the protein was transmitted to the embryo, it was not detected at the posterior pole (Figure 2B and Supplementary Figure S1E). In the presence of GFP-Vas^WT^ Aub and Ago3 showed wild-type localization in oocytes and embryos, while GFP-Vas^GNT^ only partially restored localization of the two PIWI proteins (Figure 3A-B and Supplementary Figure S2C-D and S3B-D). These observations indicate that helicase activity of Vas is necessary for stable localization of the protein itself and of Aub and Ago3 at the embryo posterior pole.

**Figure 3.**
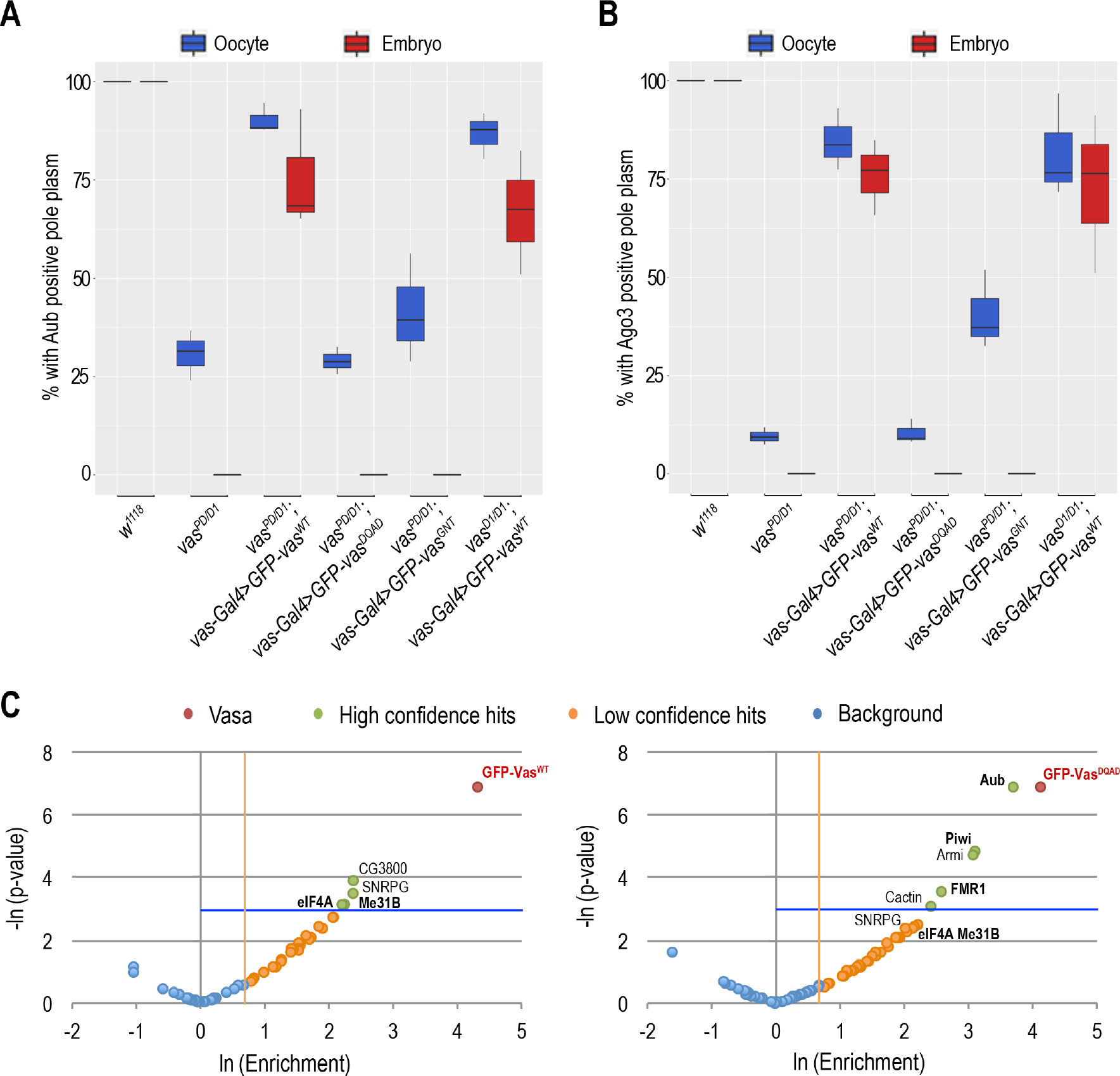
Localization of Aub and Ago3 in the egg-chamber and embryos depends on Vasa. A) Box plot showing percentage of oocytes and embryo progeny of wild-type (*w*^*1118*^*), vas*^*PD/D1*^, *vas*^*PD/D1*^*; vas-Gal4>GFP-Vas*^*WT*^, *vas*^*PD/D1*^*; vas-Gal4>GFP-Vas*^*DQAD*^, *vas*^*PD/D1*^*; vas-Gal4>GFP-Vas*^*GNT*^ *and vas*^*D1/D1*^*; vas-Gal4>GFP-Vas*^*WT*^ flies displaying Aub positive pole plasm, as determined by immunohistochemical detection of Aub. Experiments were performed in 3 independent replicates. B) Box plot representing percentage of oocytes and embryo progeny of wild-type (*w*^*1118*^*), vas*^*PD/D1*^, *vas*^*PD/D1*^*; vas-Gal4>GFP-Vas*^*WT*^, *vas*^*PD/D1*^*; vas-Gal4>GFP-Vas*^*DQAD*^, *vas*^*PD/D1*^*; vas-Gal4>GFP-Vas*^*GNT*^ *and vas*^*D1/D1*^*; vas-Gal4>GFP-Vas*^*WT*^ flies displaying Ago3 positive pole plasm, as determined by immunohistochemical detection of Ago3 protein. Experiments were performed in 3 independent replicates. C) Mass spectormetry analysis of GFP-Vas^WT^ (left panel) and GFP-Vas^DQAD^ (right panel) co-IPs. Comparison of the fold-change in abundance of proteins based on spectral count ratio (enrichment >2 fold), and statistical significance (p < 0.05) of the fold-change between GFP-Vas co-IPs and GFP negative control co-IPs. Statistically significant proteins with the highest positive fold-change were considered high confidence hits (green); proteins with a fold-change >2 but statistically non-significant (p > 0.05), were considered low confidence hits (orange); proteins below both thresholds were considered background (blue). Vasa proteins are indicated in red. Proteins previously known to interact with Vasa are in bold. Statistical analysis was performed on two biological replicates.

### Vasa associated proteins in the *Drosophila* ovary

The dynamic association of DEAD-box RNA helicases with multiprotein complexes (Linder and Jankowsky, 2011) renders challenging the biochemical detection of their interaction partners. The E400Q mutation, which locks Vas-containing protein complexes, is an ideal biochemical tool for identifying Vasa’s interaction partners *in vivo* (Xiol et al., 2014). We performed co-IP experiments from ovaries expressing GFP-Vas^WT^ and GFP-Vas^DQAD^, and identified their associated proteins by mass spectrometry. Flies expressing GFP ubiquitously served as a negative control. The quality and specificity of the coimmunoprecipitations was validated by Coomassie staining and western blot detection of several of the identified proteins (Supplementary Figure S4A-B). We identified 57 proteins associated with GFP-Vas^WT^ and 71 associated with GFP-Vas^DQAD^ (Figure 3C and Supplementary Figure S4C-E). In the case of the GFP-Vas^WT^ co-IP, the stringent conditions and absence of a cross-linking reagent (see Experimental Procedure) restricted detection to stable complexes. For instance, Oskar protein, which interacts with Vas at the posterior pole (Breitwieser et al., 1996; Jeske et al., 2017; Wang et al., 2015) was not detected in the co-IPs, whether in the case of GFP-Vas^WT^ or of the “locked” GFP-Vas^DQAD^ (Figure 2B upper panels). Among the Vas interactors we identified were Aub, Piwi, Fragile X Mental Retardation1 (FMR1) and eIF4A (Figure 3C), which have been shown to be in complex with Vas also in early embryos (Megosh et al., 2006; Thomson et al., 2008). Ago3 was not among the interactors, in agreement with previous findings that *Bombyx* Vas directly associates with Siwi (*B. mori* Aub homolog) but not with Ago3 (Nishida et al., 2015).

### Vasa activity in the germarium is essential for oogenesis

Absence of Vas in *Drosophila* females causes oogenesis arrest (Lasko and Ashburner, 1988, 1990). To determine at which stage of oogenesis Vas is required, we used either the *vas-Gal4* or the *matTub-Gal4* driver to express GFP-Vas^WT^ at distinct stages of oogenesis: the *vas* promoter is active throughout oogenesis, whereas the *matTub promoter* is inactive in the germarium, but active during the subsequent stages (Supplementary Figure S1B). *matTub-Gal4* driven expression of GFP-Vas^WT^ in *vas*^*D1/D1*^ females fully rescued oogenesis in 3-day-old flies, but as these progressed in age, oogenesis arrested (Figure 4A and Supplementary Figure S5A). In contrast, *vas-Gal4* driven expression of GFP-Vas^WT^ in the same *vas*^*D1/D1*^ background restored oogenesis independently of the age of the flies (Figure 4A and Supplementary Figure S5A). Of note, expression of helicase inactive GFP-Vas^DQAD^ and GFP-Vas^GNT^ proteins did not rescue oogenesis, whichever Gal4-driver was used. In addition, analysis of egg-chamber development showed that ovarian atrophy takes place between oogenesis stage 6 and 8 and is a result of pyknosis (Supplementary Figure 5B).

**Figure 4.**
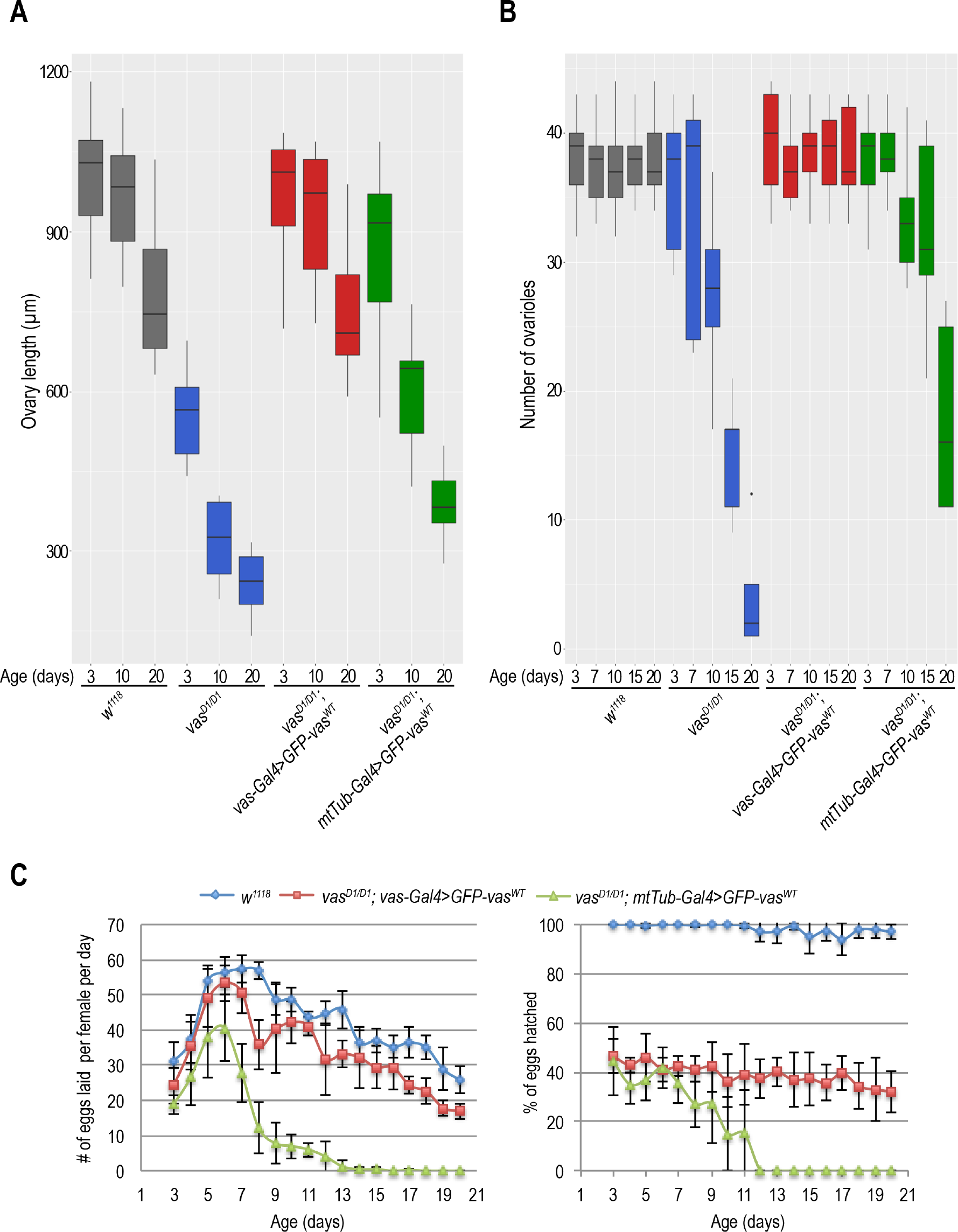
Vasa activity in the germarium is essential for germ cell development. A) Box plot representing length of ovaries of 3-day, 10-day, and 20-day old wild-type (*w*^*1118*^*), vas*^*D1/D1*^, *vas*^*D1/D1*^*; vas-Gal4>GFP-Vas*^*WT*^ and *vas*^*D1/D1*^*; matTub-Gal4>GFP-Vas*^*WT*^ flies. The measurements were performed on 10 flies (n=10). B) Box plot representing the number of egg-chamber-producing ovarioles per 3-day, 7-day, 10-day, 15-day, and 20-day old wild-type (*w*^*1118*^*), vas*^*D1/D1*^, *vas*^*D1/D1*^*; vas-Gal4>GFP-Vas*^*WT*^ and *vas*^*D1/D1*^*; matTub-Gal4>GFP-Vas*^*WT*^ female. Experiment was performed on 5 flies (n=5). C) Egg-laying rate (right) and hatching rate (left) measured daily between day 3 and day 20 post pupal enclosure of wild-type (*w*^*1118*^*), vas*^*D1/D1*^, *vas*^*D1/D1*^*; vas-Gal4>GFP-Vas*^*WT*^ and *vas*^*D1/D1*^*; matTub-Gal4>GFP-Vas*^*WT*^ females. Experiments were performed in 5 independent replicates.

**Figure 5.**
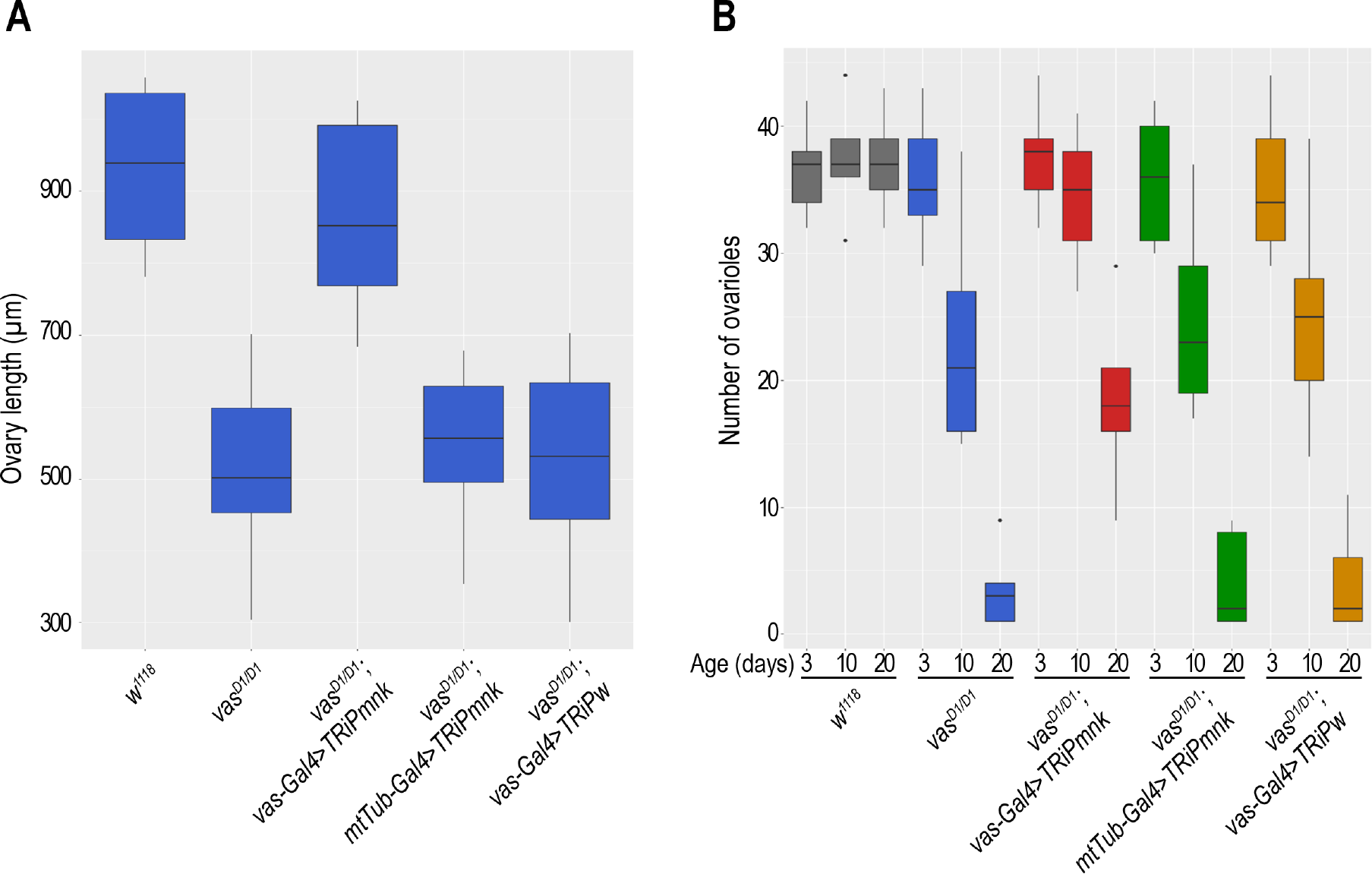
Chk2-signaling in the germarium induces arrest of germ cell development. A) Box plot representing length of ovaries of wild-type (*w*^*1118*^*), vas*^*D1/D1*^, *vas*^*D1/D1*^*; vas-Gal4>TRiPmnk*, *vas*^*D1/D1*^*; matTub-Gal4>TRiPmnk* and *vas*^*D1/D1*^*; vas-Gal4>TRiPw* flies. The measurements were performed on 15 flies (n=15). B) Box plot representing the number of egg-chamber-producing ovarioles per 3-day, 10-day and 20-day old wild-type (*w*^*1118*^*), vas*^*D1/D1*^, *vas*^*D1/D1*^*; vas-Gal4>TRiPmnk*, *vas*^*D1/D1*^*; matTub-Gal4>TRiPmnk* and *vas*^*D1/D1*^*; vas-Gal4>TRiPw* female. Experiment was performed on 5 flies (n=5).

To test whether absence of Vas in the germarium interferes with germ cell development, we determined the number of egg-chamber-producing ovarioles per female. Strikingly, in the case of *matTub-Gal4>GFP-Vas*^*WT*^ expressing *vas*^*D1/D1*^ and *vas*^*D1/D1*^; flies, the number of ovarioles decreased with the age of the females, whereas it did not, in the case of *vas-Gal4>GFP-Vas*^*WT*^ expressing *vas*^*D1/D1*^; flies or of wild-type flies (Figure 4B). Furthermore, egg-laying analysis showed that the number of eggs produced by *vas*^*D1/D1*^*; matTub-Gal4>GFP-Vas*^*WT*^ females decreased with the age of the females and eventually stopped altogether (Figure 4C left diagram). However, the hatching rate of eggs produced by *vas*^*D1/D1*^ females as a result of *matTub-Gal4* or of *vas-Gal4 GFP-Vas*^*WT*^ driven expression of *GFP-Vas*^*WT*^ did not differ significantly (Figure 4C right diagram). Taken together, these results indicate that progression and completion of oogenesis depends on the activity of Vas in the germarium.

### Chk2 signaling in the germarium induces oogenesis arrest in *vas* mutant *Drosophila*

We recently showed that *mnk* (Chk2) and *vas* interact genetically, and that depletion of Chk2 signaling in loss of function *vas*^*D1/D1*^ flies rescues oogenesis, but that the embryos die due to severe DNA damage (Durdevic et al., 2018). Moreover, previous studies determined that Vas is phosphorylated in a Chk2-dependent manner (Abdu et al., 2002; Klattenhoff et al., 2007). To investigate whether the age-dependent oogenesis arrest observed in *vas*^*D1/D1*^*; matTub-Gal4>GFP-Vas*^*WT*^ flies is due to the Chk2-signaling, we used RNAi to knock down *mnk* mRNA by either *matTub-Gal4* or *vas-Gal4* driven expression of *mnk* dsRNA (*vas*^*D1/D1*^*; matTub-Gal4>TRiPmnk and vas*^*D1/D1*^*; vas-Gal4>TRiPmnk*, respectively). Knock-down efficiency tests using quantitative PCR showed *mnk* mRNA levels to be between 30% and 40% of the wild-type *mnk* level (Supplementary Figure S6A). However, fluorescent *in situ* RNA hybridization (FISH) analysis showed that upon *matTub-Gal4* driven knockdown of *mnk*, the mRNA was detectable in the germarium and not in the later stages of oogenesis, whereas *vas-Gal4* driven *mnk*-RNAi downregulated *mnk* throughout oogenesis (Supplementary Figure S6B). Furthermore, whereas *vas-Gal4* driven silencing of *mnk* in *vas*^*D1/D1*^ females restored oogenesis, *matTub-Gal4* driven knockdown of *mnk* did not (Figure 5A). This indicates Chk2-mediated signaling activity in the germarium determines the fate of developing egg-chambers. Although the efficiency of *vas-Gal4* RNAi-driven downregulation of *mnk* decreased over time, finally resulting in ovarian atrophy, we observed a more severe age-dependent decrease in the number of egg-chamber-producing ovarioles when *mnk* knockdown was driven by *matTub-Gal4* (Figure 5B). These results suggest that in *vas* mutants Chk2-signaling specifically in the germarium induces arrest of germ cell development.

## Discussion

We have demonstrated that development of the *Drosophila* female germline depends on Vas activity in early oogenesis. Our data indicate that progression and completion of oogenesis require helicase active Vas. However, as our fusion proteins show low expression levels, we cannot rule out that when expressed at higher levels Vas might support oogenesis independently of helicase activity (Dehghani and Lasko, 2015). Using stage-specific promoters, we manipulated the expression of Vas and determined that activity of Vas in the germarium is crucial for germ cell lineage development. Our conclusion that oogenesis depends on an early helicase activity of Vas is consistent with the finding that Vas directly interacts with *meiotic P26* (*mei-P26)* mRNA and activates its translation (Liu et al., 2009). Mei-P26 itself has been found to cooperate with proteins such as Bag of marbles and Sex lethal to promote both GSC self-renewal and germline differentiation (Li et al., 2012; Li et al., 2013). In addition, we identified Vas interaction partners Lingerer, Rasputin, FMR1 and Caprin, which have been shown to cooperate in restricting tissue growth in a non-germline tissue, the *Drosophila* eye (Baumgartner et al., 2013). Interestingly, these proteins were found to interact in *Drosophila* ovaries as well (Costa et al., 2013; Costa et al., 2005) suggesting formation of a complex that could act in conjunction with Vas to control growth of *Drosophila* germline tissue. FMR1 was previously shown to interact with Vas in embryos and to be important for PGC formation (Deshpande et al., 2006; Megosh et al., 2006). In *Drosophila* ovaries, FMR1 was proposed to participate in the regulation of germline proliferation and GSC maintenance (Epstein et al., 2009; Yang et al., 2007). In addition, Vas interaction with numerous other factors implicated in promoting the self-renewal of GSC such as Rm62 (Ma et al., 2017), Bel (Kotov et al., 2016), and Nop60B (Kaufmann et al., 2003) indicates an intricate network of Vas-associated processes involved in sustaining the germ cell lineage. Further studies will be required to determine how these proteins collaborate to regulate early germ cell development.

In *Drosophila,* Chk2-signaling triggered by DNA damage, replication stress or nuclear lamina dysfunction induces GSC loss (Barton et al., 2018; Ma et al., 2016; Molla-Herman et al., 2015). Our previous study showed that removal of Chk2 in *vas* mutant flies fully restores oogenesis, while progeny embryos succumb to transposon up-regulation and DNA damage (Durdevic et al., 2018). Here we went further and genetically determined that, in *vas* mutants, *Drosophila* oogenesis arrests due to Chk2-signaling in the germarium. Furthermore, the decline of ovariole number is Chk2-dependent, indicating that in *vas* mutants Chk2-signaling compromises germ cell lineage development. An earlier study suggested that Vas interacts with Aub to regulate mitotic chromosome condensation in *Drosophila* GSCs and that consequently *vas* mutants display aberrant chromosome segregation during GSC mitosis (Pek and Kai, 2011). Defects in mitotic chromosome segregation can be the cause as well as the consequence of Chk2 activation in the respective daughter cells (Bakhoum et al., 2014; Jansen et al., 2011). As *vas* mutant GSCs do not display DNA damage (Pek and Kai, 2011), we speculate that mitotic chromosome segregation defects trigger Chk2-signaling in the daughter GSC and the CB, disrupting GSC self-renewal and development of the germ cell lineage. We conclude that the conserved RNA helicase Vas plays an essential role in sustaining the germ cell lineage in *Drosophila*.

## Data availability

The authors declare that all data supporting the findings of this study are available within the manuscript and its supplementary files.

## Author contributions

Z.D. and A.E. conceived and designed the experiments. Z.D carried out the experiments and analysed the data. Z.D. and A.E. wrote the manuscript.

## Acknowledgments

We thank Paul Lasko for sharing his insight into Vasa helicase function. We thank Mikiko Siomi for the gift of antibodies against Ago3, Beat Suter for the pUAS-attB transgenesis vector, and Hugo Bellen, Jean-René Huynh and Stefano De Renzis for fly stocks. We thank the EMBL Proteomics and Advanced Light Microscopy Core Facilities. We are grateful to Anna Cyrklaff and Alessandra Reversi for their help with experiments and to Sandra Mueller for *Drosophila* transgenesis. This work was funded by the EMBL and Z.D. by a postdoctoral fellowship from the EMBL Interdisciplinary Postdoc Program (EIPOD) under Marie Curie COFUND actions.

